# Beta-cell Cre expression and reduced *Ins1* gene dosage protect mice from type 1 diabetes

**DOI:** 10.1101/2022.02.03.479027

**Authors:** Søs Skovsø, Peter Overby, Jasmine Memar-Zadeh, Jason T.C. Lee, Jenny C.C. Yang, Iryna Shanina, Vaibhav Sidarala, Elena Levi-D’Ancona, Jie Zhu, Scott A. Soleimanpour, Marc S. Horwitz, James D. Johnson

## Abstract

A central goal of physiological research is the understanding of cell-specific roles of disease-associated genes. Cre-mediated recombineering is the tool of choice for cell type-specific analysis of gene function in pre-clinical models. In the type 1 diabetes research field, multiple lines of NOD mice have been engineered to express Cre recombinase in pancreatic β-cells using insulin promoter fragments, but tissue promiscuity remains a concern. Constitutive *Ins1*^tm1.1(cre)Thor^ (*Ins1*^Cre^) mice on the C57/bl6-J background has high β-cell specificity and with no reported off-target effects. We explored if NOD:*Ins1*^Cre^ mice could be used to investigate β-cell gene deletion in type 1 diabetes disease modeling. We studied wildtype (*Ins1*^WT/WT^), *Ins1* heterozygous (*Ins1*^Cre/WT^ or *Ins1*^Neo/WT^), and *Ins1* null (*Ins1*^Cre/Neo^) littermates on a NOD background. Female *Ins1*^Neo/WT^ mice exhibited significant protection from diabetes, with further near-complete protection in *Ins1*^Cre/WT^ mice. The effects of combined neomycin and Cre knock-in in *Ins1*^Neo/Cre^ mice were not additive to the Cre knock-in alone. In *Ins1*^Neo/Cre^ mice, protection from diabetes was associated with reduced insulitis at 12 weeks of age. Collectively, these data confirm previous reports that loss of *Ins1* alleles protects NOD mice from diabetes development and demonstrates, for the first time, that Cre itself may have additional protective effects. This has significant implications for the experimental design and interpretation of pre-clinical type 1 diabetes studies using β-cell-specific Cre in NOD mice.

## Introduction

Type 1 diabetes (T1D) is a chronic disorder precipitated by immune-mediated pancreatic β-cell destruction and associated with the presence of autoantibodies against β-cell proteins (1,2). Due to the progressive loss of β-cells and consequent insulin deficiency, individuals living with type 1 diabetes have lifelong dependency on exogenous insulin (3). Higher levels of endogenous insulin secretion are associated with better short-and long-term outcomes in type 1 diabetes. Preservation of residual β-cells and islet function at the time of diagnosis is therefore imperative (4).

Exogenous insulin administration is not a cure of type 1 diabetes. Research efforts using pre-clinical animal models continue to produce new therapeutic possibilities. Initially developed as a model for spontaneous onset of cataracts, the female non-obese diabetic (NOD/ShiLt) mouse (commonly known as NOD) is the most well-established and extensively used model of type 1 diabetes (5). The polygenic NOD mouse model recapitulates multiple pathophysiological features of human type 1 diabetes, including the development of autoantibodies in prediabetic NOD mice (6), circulating autoreactive T cells (7), and subsequent onset of hyperglycemia as progressive β-cell loss materializes (8). While the emergence of hyperglycemia most often occurs between 12-15 weeks of age, islet immune infiltration is established earlier at 8-12 weeks with insulitis present across the pancreas(9). The cleanliness of housing facilities and the animal breeding approach are just two factors which affect the age at which these features present.

Mice express two nonallelic insulin genes. The insulin 2 gene (*Ins2*) is the murine homologue of the human insulin gene and is located on chromosome 7 (10). The insulin 1 gene (*Ins1*) is the result of an RNA-mediated gene duplication event. *Ins1* has a simpler gene structure lacking the second intron present in *Ins2*, and is found on the murine chromosome 19 (10). *Ins1* is expressed specifically in pancreatic β-cells, whereas *Ins2* is expressed predominantly in β-cells, with trace expression in other tissues including thymus and brain (11). Previous studies have examined the effects of insulin gene knockout in NOD mice. Complete knockout of *Ins2* on the NOD background accelerated diabetes onset (2), a phenomenon attributed to a failure in central tolerization to insulin. Thymus-specific deletion of *Ins2* was reported to be sufficient to cause spontaneous diabetes, even outside the NOD background (12). Although we have not observed autoimmune diabetes in globally deficient *Ins2*^-/-^ mice (13,14) it has previously reported that male, but not female, NOD:*Ins1*^Neo/WT^;*Ins2*^-/-^ mice, with a single remaining *Ins1* allele have been shown to succumb to insulin insufficiency (15), a finding we confirmed on other backgrounds (16). These observations were found to be dependent on housing conditions (11). In contrast, NOD:*Ins1*^Neo/Neo^ mice have previously been shown to be largely protected from insulitis diabetes (10), which was proposed to be due to the loss of autoantigenic *Ins1*-derived peptides despite presence of insulin autoantibodies (IAA). Insulin is a primary auto-antigen in both murine and human type 1 diabetes pathogenesis (1). Replacing *Ins1* with non-antigenic human insulin also protects NOD mice from onset of diabetes (17). Together, these previous observations indicate that insulin 1 gene dosage is a key driver of diabetes in NOD mice.

To advance the understanding of the underlying mechanisms of T1D pathogenesis, powerful genetic engineering tools are employed by researchers across the world. *Ins1*^Cre^ mice take advantage of the tissue specificity of *Ins1* expression to excise *lox*P site-flanked (floxed) DNA segments. As the Cre DNA recombinase allele is inserted into exon 2 of *Ins1* in these mice, floxed target genes are removed specifically in beta-cells at the cost of the loss of 1 *Ins1* allele. Both *Ins1*^Cre^ and *Ins1*^Neo^ mice have reduced *Ins1* gene dosages with 50% or 100% in their heterozygeous and homozygeous states, respectively. For *Ins1*^Neo^ mice, the neomycin cassette obliterates the entire *Ins1* promotor and gene sequence in addition to deletion of 7-9kb upstream and downstream of *Ins1* (see Figure 1A).

**Figure 1.**
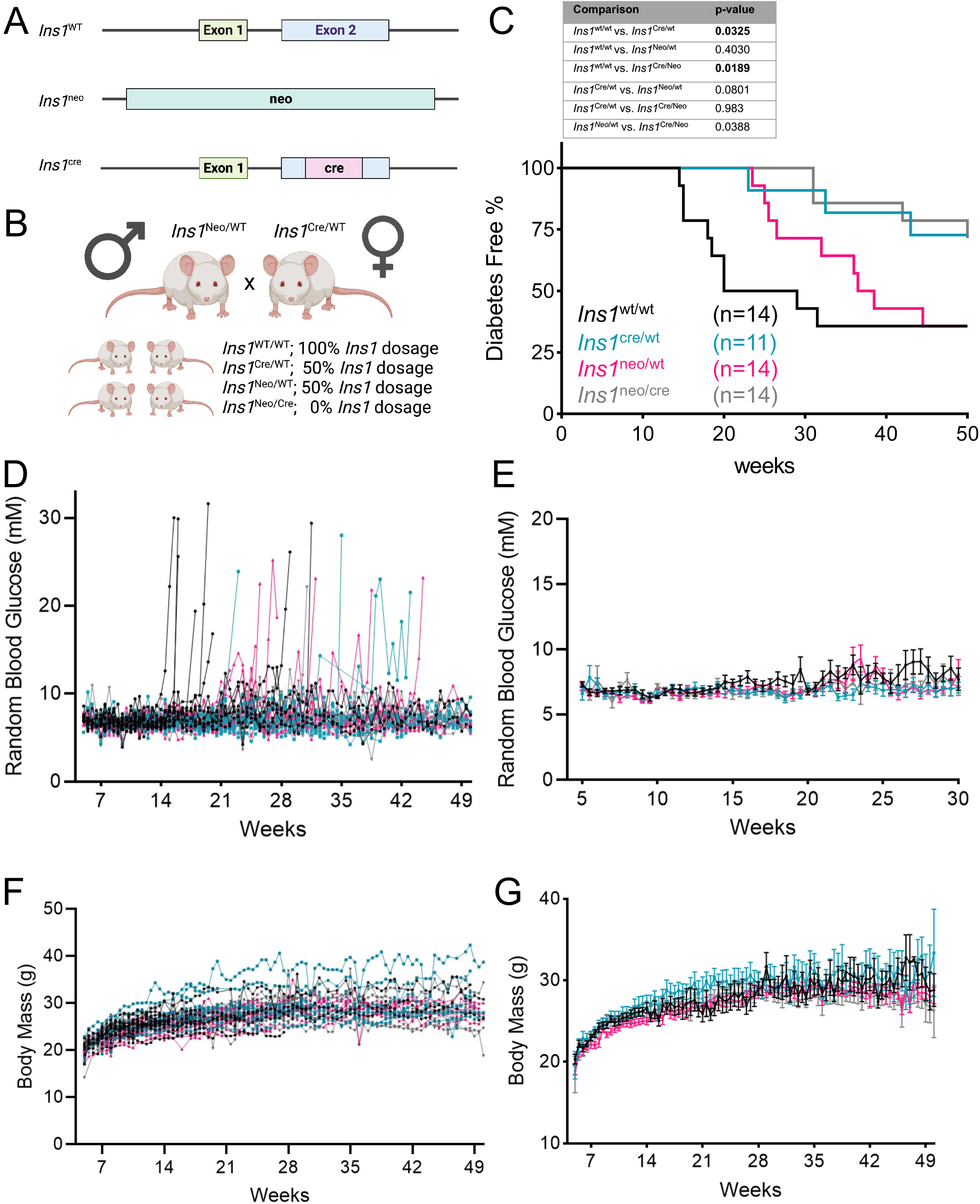
*Ins1* replacement with Cre and Neo protects female NOD mice from type 1 diabetes. **(A)** Structure of the wildtype (WT) *Ins1* locus, recombinant alleles result of the neomycin (neo) targeting vector, or the *Ins1* Cre locus. Created with Biorender.com. **(B)** Overview of breeding strategy. Created with Biorender.com. **(C)** Kaplan-Meier plot denoting diabetes incidence in NOD mice by *Ins1* genotype, including *Ins1*^WT/WT^, *Ins1*^Cre/WT^, *Ins1*^Neo/WT^ and *Ins1*^Neo/Cre^. Survival analysis was performed using Log-rank (Mantel-Cox) test (*p* value shown). **(D**,**E)** Individual and mean random blood glucose of female mice. The mean blood glucose of the *Ins1*^Neo/Cre^ was significantly lower than that of the *Ins1*^WT/WT^ colonies, with an adjusted p-value of 0.0023. Mean blood glucose of the *Ins1*^Neo/WT^ colony was higher than those of the *Ins1*^WT/WT^ and *Ins1*^Cre/WT^ colonies, with adjusted p-values of 0.0002 and 0.0058 respectively. **(F**,**G)** Individual and mean body mass traces female mice. The mean body mass of the *Ins1*^Neo/Cre^ mice was significantly lower compared to the *Ins1*^Cre/WT^ colony (adjusted p-value<0.0001). The mean blood glucose of the *Ins1*^Cre/WT^ was also higher than the *Ins1*^WT/WT^ and the *Ins1*^Cre/WT^ colonies, both with an adjusted p-value<0.0001. Error bars represent SEM

Since the early days of Cre/*lox*P system usage, concerns of side effects such as Cre toxicity have been raised. More than two decades ago, Loonstra *et al* observed increased sister chromatid exchange frequency, leading to a cellular halt in the G_2_/M phase and a reduction in proliferation of mouse embryonic fibroblasts (MEF) transduced with a bicitronic retroviral vector encoding Cre (18). Silver and Livingston demonstrated similar results including ceased proliferation in 293xLac cells, NIH 3T3 cells and MEFs transfected with Cre-expressing retroviral vectors (19). The same study revealed chromosomal abnormalities in Cre-expressing MEFs. Lentiviral Cre administration and expression have likewise been shown to reduce proliferation through cellular accumulation in the G_2_M phase of Cre-expressing CV-1 and COS cells, two kidney cell lines derived from monkeys (20). Cre toxicity has been shown to be dose dependent, even when expressed transiently, when using an adenovirus vector (21). Past studies have confirmed Cre toxicity and apoptosis in mouse cardiac tissue (22,23) and p53^-/-^ thymic lymphoma cells (24). Collectively, these findings emphasize the necessity of using Cre-only (no flox) controls into *in vivo* research study designs since Cre recombinase expression may affect cells beyond the intended target gene deletion.

For this study, we generated and characterized a new NOD *Ins1*^Cre^ knock-in mouse line. Our intension was to establish an *in vivo* tool facilitating the unravelling of type 1 diabetes pathogenesis. The NOD *Ins1*^Cre^ model would ideally enable studying of β-cell specific deletion of target genes in mice on an NOD background. As a predecessor, the goal of this study was to investigate the impact of 1) deleting one *Ins1* allele and 2) introducing beta cell specific Cre expression, on the spontaneous diabetes development NOD mice are well known for. As previous studies by the Eisenbarth group provided evidence that *Ins1*^Neo/WT^ mice, with only one intact *Ins1* allele, exhibit decreased and delayed diabetes incidence, we were expecting similar results for *Ins1*^Cre/WT^ mice. We incorporated both NOD:*Ins1*^Neo/Wt^ NOD;*Ins1*^WtTWT^ littermates in our study design, to control for both the loss of one *Ins1* allele and introduction of Cre expression in NOD *Ins1*^Cre/WT^ mice. We compared diabetes incidence, insulitis severity, and immune activation between groups within each sex. Our data demonstrate that *Ins1*-driven Cre expression has further protective effects beyond the loss of a single *Ins1* allele deleted via *Ins1*^*Neo*^. Carefully chosen proper controls and caution are therefore required when interpreting experiments including β-cell specific Cre expression in mice on an NOD background.

## Methods

### Mice

All animal procedures and ethical standards were in accordance with the Canadian Council for Animal Care guidelines. All animal studies and protocols were approved by the University of British Columbia Animal Care Committee and Institutional Care and Use Committee (IACUC) at the University of Michigan. At UBC, mice were housed in the Centre for Disease Modelling Specific Pathogen-Free (SFP) facility on a standard 12-h light/12-h dark cycle with ad libitum access to chow diet (PicoLab, Mouse Diet 20–5058). To generate NOD:*Ins1*^Cre^ and NOD:*Ins1*^Neo^ mice, we contracted The Jackson Laboratories to backcross (>12 times) B6(Cg)-*Ins1*^*tm1*.*1(cre)Thor*^/J (The Jackson Laboratory, US, #026801) (*Ins1*^Cre^) and NOD.129S2(B6)-*Ins1*^*tm1Jja*^/GseJ (The Jackson Laboratory, US, #0005035) *Ins1*^Neo^) mice onto a NOD/ShiLtJ (The Jackson Laboratory, US, #001976) (NOD) background. The *Ins1*^Cre^ and *Ins1*^Neo^ mice were originally on a mixed, largely C57Bl/6J, background. Subsequently, we designed a strict breeding strategy for our study. An *Ins1*^Cre^ maternal parent colony as well as a *Ins1*^Neo^ paternal parent colony was established (Fig. 1B). Each parent colony was backcrossed every five generations. Female *Ins1*^Cre/WT^ and male *Ins1*^Neo/WT^ mice were set up as breeders, at 7-8 weeks of age, to generate experimental mice. Mice from the parental colonies were only included once as breeders to eliminate risk of onset of hyperglycemia during pregnancy and weaning at later ages. This breeding strategy generated littermates of four genotypes: wildtype NOD:*Ins1*^WT/WT^ mice with both insulin 1 alleles, heterozygous NOD:*Ins1*^Cre/WT^ mice with an *Ins1* replaced with Cre-recombinase, heterozygous NOD:*Ins1*^Neo/WT^ mice with one *Ins1* allele replaced with a neomycin cassette, and full *Ins1* null mice with both *Ins1* alleles replaced NOD:*Ins1*^Neo/Cre^. Specific cohorts were monitored twice per week for the sole purpose of determining hyperglycemia onset incidence and body mass changes. Any mice that developed diabetes, defined as two consecutive blood glucose measurements ≥16 mmol/L or one measurement ≥22 mmol/L, were euthanized. Specific cohorts were generated for tissue analysis terminated at 12 weeks of age in a pre-diabetic phase (for the Vancouver housing facility) or at 1 year of age, all animals were monitored for diabetes prior to euthanasia.

An additional colony of NOD:*Ins1*^Cre^ mice were generated independently through in-house backcrossing at a second site (University of Michigan) via the speed congenic approach in consultation with Charles River Laboratories (Wilmington, MA). Following each backcross, *Ins1*^Cre/WT^ offspring with allelic profiles most closely matching the NOD strain (determined by MAX-BAX mouse 384 SNP panel screening), were selected as breeders for the subsequent backcross. Following 8 generations of backcrossing, animals with an allelic profile percent match >99.9% were utilized to generate *Ins1*^WT/WT^ and *Ins1*^Cre/WT^ experimental mice. Animals were housed in an SPF facility on a standard 12-h light/12-h dark cycle with access to ad libitum chow diet (LabDiets, Rodent Diet 5L0D) and water. Drinking water was provided at a pH level of 2.5-3 upon advice from Jackson laboratories that acidified water supports the diabetes frequency in many NOD colonies (25)Blood glucose measurements were taken once-twice per week. Incidence of diabetes, defined as blood glucose levels > 16 mM for 5 consecutive measurements, was recorded. Diabetic mice were euthanized and excluded from future analysis.

### Tissue processing and histology

Immediately following euthanasia, pancreata were collected according to a pre-established protocol with the exception that extracted pancreases were not further treated to obtain isolated islets and were instead processed as a whole(26). The dissected pancreata were fixed in 4% PFA for 24 hours prior to storage in 70% ethanol at 4 °C. Paraffin-embedded sections were prepared, stained with hematoxylin and eosin (H&E), and imaged by WaxIT Histology Services Inc. (Vancouver, BC).

### Islet infiltration scoring

Images from H&E-stained pancreatic sections were analyzed with QuPath software (27). Islets in pancreatic sections were scored blindly for pancreatic islet infiltration of mononuclear immune cells according to the previously established 4-point scale(28). In brief, 0-no insulitis, 1-peri-insulitis marked by less than 25% peripheral immune-islet infiltration, 2-insulitis marked by 25-75% immune cell infiltration, 3-severe insulitis marked by greater than 75% immune-islet infiltration. Random samples were scored by a second person to ensure consistency. 25-30 islets per mouse.

### Insulin auto-antibodies and plasma insulin

Blood for plasma insulin and autoantibodies measurements was collected at study endpoint by cardiac puncture from anesthetized Animals (2.5% isoflurane) and put directly on ice. Insulin plasma where measured using Mouse Insulin ELISA kit (#80-INSMS-E10, RRID: AB_2923075; ALPCO). Samples were shipped to the Insulin Antibody Core Lab at the University of Colorado, Barbara Davis Diabetes Centre for auto-antibody analysis. The IAA data are expressed as an index against a standard positive and negative controls as procedure at the Insulin Antibody Core lab

### Flow cytometry

Immediately following euthanasia lymph nodes and spleen were isolated and splenic single-cell suspensions were counted and stained with fluorescently conjugated monoclonal antibodies (mAbs) for cell-surface markers (see Table 1 for list of antibodies used). Fixable Viability Dye eFluor™; 506 was used as viability dye (#65-0866-14; ThermoFischer). Following staining, cells were analyzed by flow cytometry and Flow Jo software (Tree Star Inc., Ashland, Oregon).

**Table 1.** Summary of Antibodies used for Flow cytometry.

| Antibody | Manufacturer | Colour | Laser (nM) | Filter | Catalogue# | RRID |
| --- | --- | --- | --- | --- | --- | --- |
| CD3 | ThermoFisher | e-Flour 540 | 405 | 450/45 | 48-003382 | AB_2016704 |
| CD4 | ThermoFisher | BV650 | 405 | 660/10 | 64-0042-82 | AB_2662401 |
| CD8 | BioLegend | PE-TR | 561 | 610/20 | 100762 | AB_2564027 |
| CD11c | ThermoFisher | PE | 561 | 585/42 | 12-0114-81 | AB_465551 |
| CD19 | ThermoFisher | SB780 | 405 | 763/43 | 78-0193-82 | AB_2722936 |
| CD44 | ThermoFisher | APC | 633 | 660/10 | 17-0441-82 | AB_469390 |
| CD62L | ThermoFisher | PerCP-Cy5.5 | 488 | 690/50 | 45-0621-82 | AB_996667 |
| CD69 | ThermoFisher | FITC | 488 | 525/40 | 11-0692-82 | AB_11069282 |
| Foxp3 | ThermoFisher | AF-700 | 633 | 712/25 | 565773-82 | AB_1210557 |

### Statistics

Statistical significance was assessed using 2-way ANOVA analysis with Tukey’s multiple comparisons test, at a threshold of p < 0.05. Plasma insulin, Insulitis and insulin auto-antibodies significance were quantified by 1-way ANNOVA analyses. We used the Mantel-Cox-log rank test, corrected with the Benjamini-Hochberg procedure, to analyse the Kaplain Meier survival plots. Repeated measured Mixed effect models were applied for RBG and Body mass analyses.

Prism 9 (GraphPad Software Inc., USA) was used for statistical analyses and generation of most figure panels. Data are expressed as mean ± SEM unless otherwise specified.

## Results

### Type 1 diabetes incidence

A greater proportion of female, in comparison to male, mice develop diabetes and the timing of the development of the disease is more consistent. We observed the highest diabetes incidence in female NOD:*Ins1*^WT/WT^ mice, with 65% diabetes incidence at 1 year (Fig. 1C). The majority of these mice (7/9) were diabetic before 20 weeks of age. The diabetes incidence and disease time-course of this study was comparable with previous NOD:*Ins1*^WT/WT^ cohorts in our facilities (28), although slightly delayed relative to some other facilities (see below). Female NOD:*Ins1*^Cre/WT^ exhibited both a delay in diabetes onset and a diabetes incidence by the end of the study of only 27%, which was significantly different when compared to NOD:*Ins1*^WT/WT^ mice at one year of age (p=0.033; p_adjust_=0.078; Fig. 1C). We also observed at delay in the timing of the onset between female NOD:*Ins1*^Neo/WT^ mice (Fig. 1C), in agreement with previous studies of NOD:*Ins1*^Neo/WT^ mice (10), but there was no difference in the final diabetes incidence between female littermate NOD:*Ins1*^Neo/WT^ mice (64%) and NOD:*Ins1*^WT/WT^ mice (64%) (p=0.40; p_adjust_=0.48; Fig. 1C). Double mutant female NOD:*Ins1*^Neo/Cre^ mice (lacking both wildtype alleles of *Ins1*) were protected from diabetes (29% diabetes incidence) when compared to NOD:*Ins1*^WT/WT^ mice (64%), (p=0.019; p_adjust_=0.078), to a similarly extent as NOD:*Ins1*^Cre/WT^ (27%) mice (p=0.98; p_adjust_=0.98; Fig. 1C). NOD:*Ins1*^Neo/WT^ mice had a significantly lower diabetes incidence than NOD:*Ins1*^Cre/Neo^ mice (p=0.038; p_adjust_=0.078; Fig. 1C) further suggesting Cre expression rather than loss of one *Ins1* allele protects against diabetes onset in NOD mice. No differences were observed in random blood glucose (Fig. 1D,E) and body mass (Fig. 1F,G) prior to diabetes onset in female mice. Together, these data confirm previous findings that reduced *Ins1* gene dosage protects NOD mice from diabetes, but also reveal a further, additive protective effect of β-cell Cre expression.

Male NOD:*Ins1*^WT/WT^ mice demonstrated a low diabetes incidence as expected (20% incidence), We observed no cases of hyperglycemia in any of the male mice with reduced *Ins1* gene dosage (Fig. 2A). There were no differences in random blood glucose (Fig. 2 B,C) or body mass (Fig. 2D,E) between any of the groups. These results confirm the strong sex bias in this NOD model and suggest that the protection conferred by reducing *Ins1* gene dosage may not be sex-specific.

**Figure 2.**
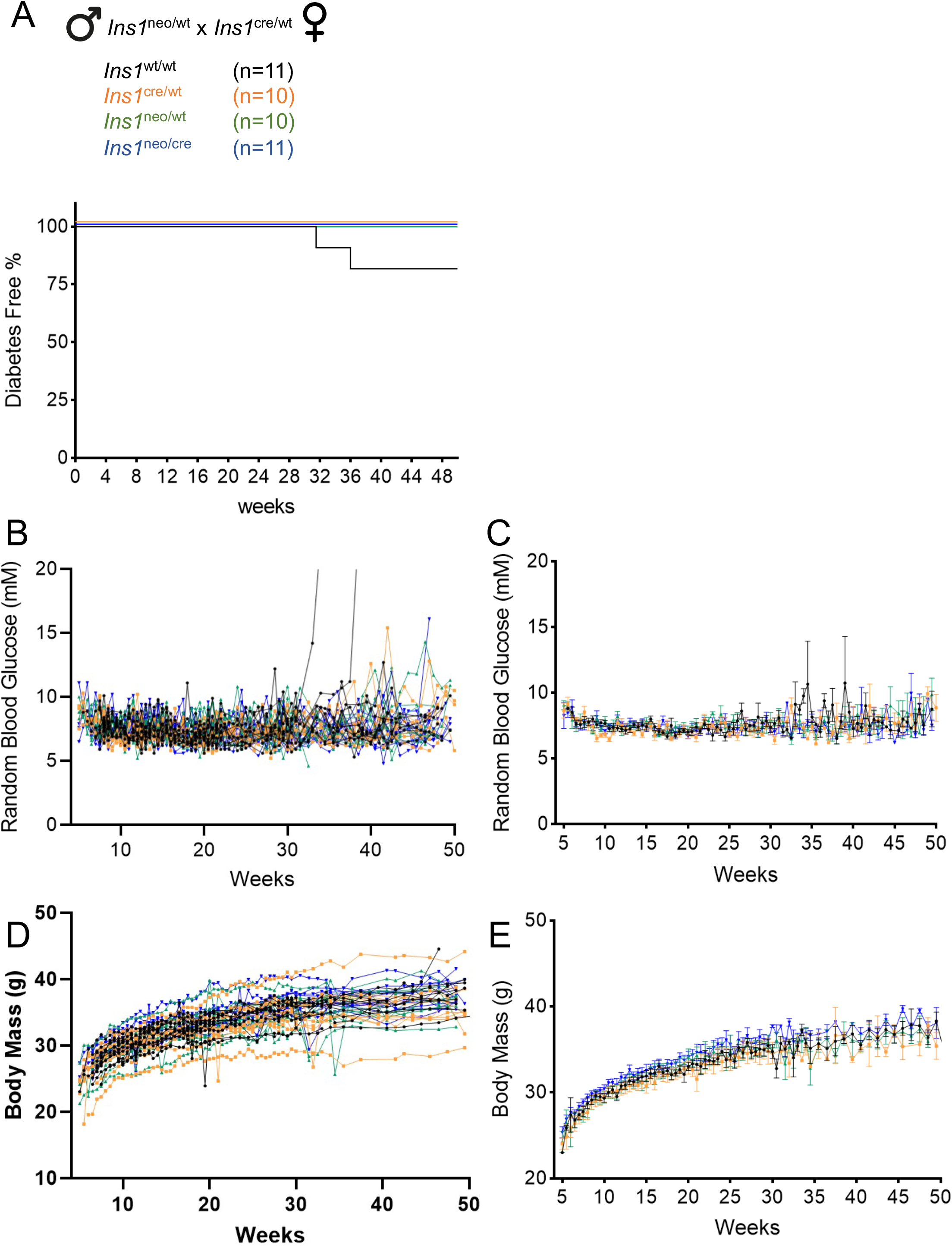
Effects of *Ins1* replacement with Cre and Neo in male NOD mice. **(A)** Kaplan-Meier plot denoting diabetes incidence in NOD mice by *Ins1* genotype. **(B**,**C)** Individual and mean random blood glucose in male mice. **(D**,**E)** Individual and mean body mass traces in male mice. Error bars represent SEM.

### Insulitis and Insulin auto-antibodies

Next, we examined the effects of reduced *Ins1* gene dosage and Cre expression on insulitis in female mice, the pathological evidence of islet directed autoimmunity. We found no significant difference in plasma insulin or autoantibodies between any of the genotypes (Fig. 3A,B). H&E-stained pancreas sections (Fig. 3C) were blindly scored for immune islet infiltration in a pre-diabetic cohort of littermates euthanized at 12 weeks of age, and also in the mice that survived to 1 year. Insulitis scores from 12-week-old mice with reduced *Ins1* gene dosages were not significantly different from NOD:*Ins1*^WT/WT^ littermates. We noticed the least amount of immune islet infiltration in double mutant NOD:*Ins1*^Neo/Cre^ mice (Fig. 3D), consistent with their more complete protection from diabetes. The outcomes of the samples gathered at 1 year are unfortunately inconclusive due to low sample size (NOD:*Ins1*^WT/WT^, n=3) (Fig 3E). The lower survival rate may therefore mask any potential significant changes for both insulitis scoring (Fig. 3E) and insulin auto-antibodies (Fig. 3B). Unfortunately, we did not collect blood for this analysis at the 12-week timepoint for all cohorts due to challenges brought about by the COVID19-pandemic.

**Figure 3.**
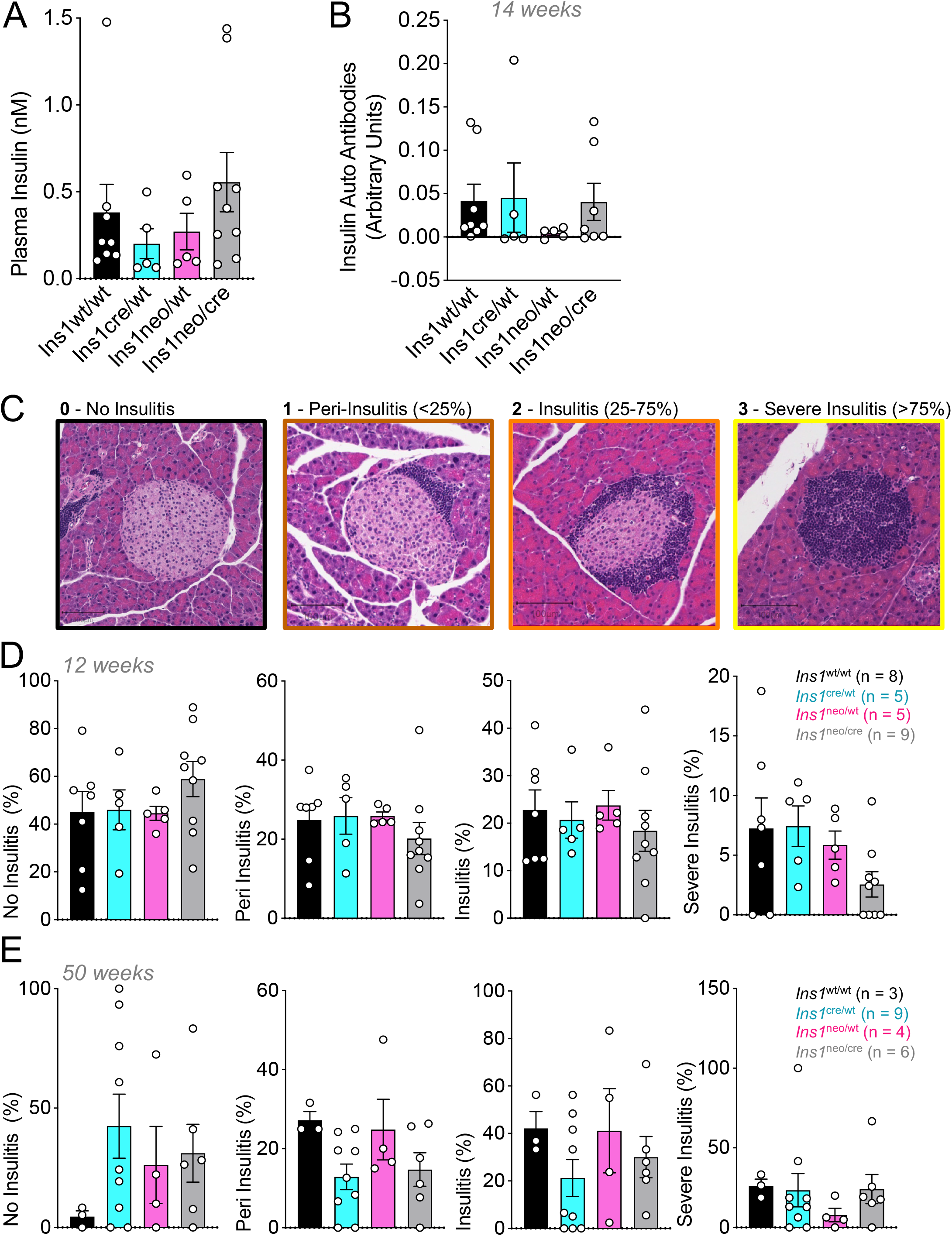
Insulitis scoring in female NOD mice with *Ins1* replacement. **(A)** Plasma insulin concentration Nanomolar (nM) collected at 14 weeks by cardiac puncture. **(B)** Mouse insulin autoantibodies collected at 14 weeks by cardiac puncture, result of IAA is expressed as an index, against internal standard positive and negative controls. **(C)** Representative images of H&E stained pancreata used for insulitis scoring. Scale bars are 100 μm. **(D)** Mean percent insulitis scores at 12-week-old of each genotype, categorized by score, 0 - No Insulitis, 1 - Peri-Insulitis (<25%), 2 - Insulitis (25-75%) and 3 - Severe Insulitis (>75%) respectively **(E)** 1-year-old time points. Error bars represent SEM.

### Immune cell characterization

To assess immune cell populations in the pancreatic lymph nodes and spleen at 50 weeks of age, we used a panel of validated antibodies for flow cytometry. While we were able to confidently identify many key immune cell populations (Fig. 4), there were no significant differences between groups (Fig. 5). These observations demonstrate that β-cell specific insulin gene manipulations alter type 1 diabetes incidence without robust effects on the lymphocytes found in the pancreatic lymph nodes and spleen.

**Figure 4.**
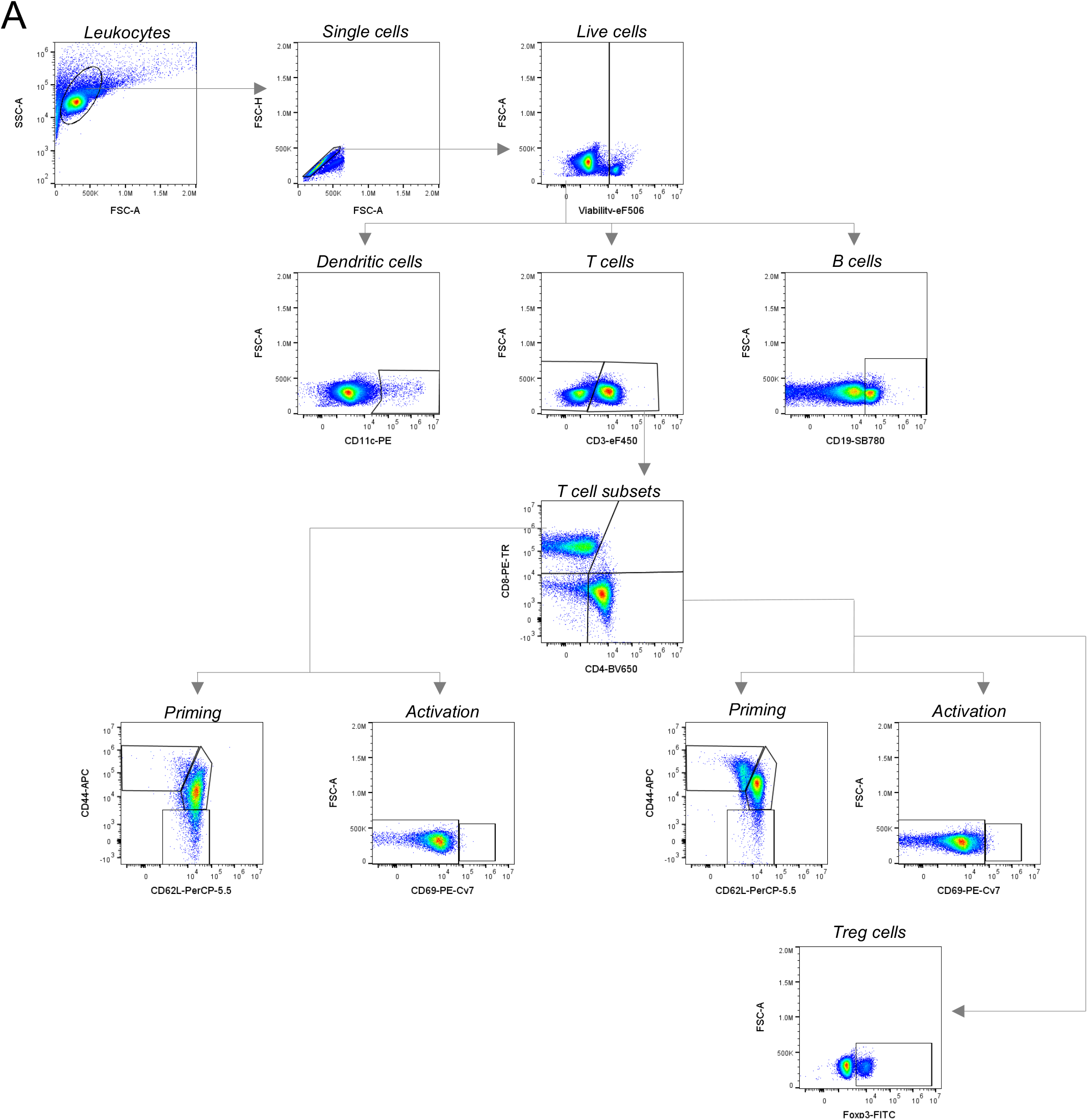
Gating strategy for flow cytometry of pancreatic lymph node and spleen cells. **(A)** Singlets were obtained with use of FSC-A x FSC-H parameters and viable cells were identified by selecting viability dye negative cells for subsequent analysis. The populations were subsequently split into three groups of interest with dendritic cells identified by CD11c, B cells identified by CD19, and T cells identified by CD3. T cells were further categorized into cytotoxic and helper phenotypes with use of CD4 and CD8 markers and their respective single marker populations were assessed for activation and priming status (naïve, effector, memory) with the use of CD69, CD44, and CD62L. Regulatory T cell populations were further selected for with the use of a Foxp3 marker.

**Figure 5.**
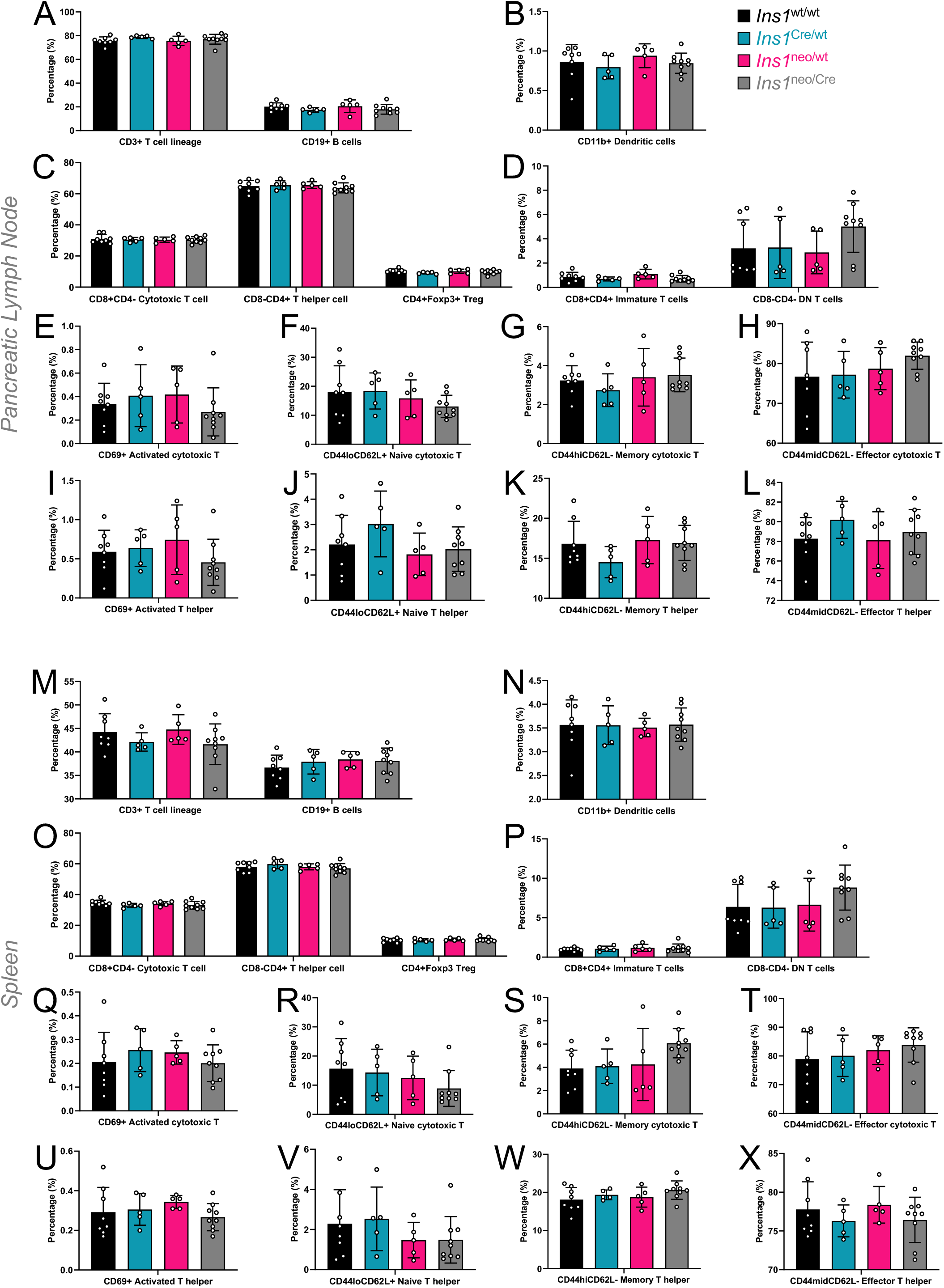
Immune profiling in female NOD mice with *Ins1* replacement. Flow cytometric analysis of cell populations within the pancreatic lymph node and spleen at 50 weeks of age. **(A)** Percentage of CD3+ positive T cells and CD19+ B cells from the pancreatic lymph node. **(B)** Percentage of CD11+ Dendritic cells from the pancreatic lymph node. **(C)** Percentage of CD8+CD4-Cytotoxic T cells, CD8+CD4-T helper cells and CD4+Foxp3+ Treg cells from the pancreatic lymph node. **(D)** Percentage of CD8+CD4+ immature cells and CD8-CD4-DN T cells from the pancreatic lymph node. **(E)** Percentage of CD69+ Activated cytotoxic T cells from the pancreatic lymph node. **(F)** CD44loCD62L + Naive cytotoxic T cells from the pancreatic lymph node. **(G)** Percentage of CD44hiCD62 – Memory cytotoxic T cells from the pancreatic lymph node. **(H)** Percentage of CD44midCD62 – Effector cytotoxic T cells from the pancreatic lymph node. **(I)** Percentage of CD69+ Activated T helper cells from the pancreatic lymph node. **(J)** CD44loCD62L + Naive T helper cells from the pancreatic lymph node. **(K)** Percentage of CD44hiCD62 + Memory T helper cells from the pancreatic lymph node. **(L)** Percentage of CD44midCD62 – Effector T helper cells from the pancreatic lymph node. **(M)** Percentage of CD3+ positive T cells and CD19+ B cells from the Spleen. **(N)** Percentage of CD11+ Dendritic cells from the spleen. (**O)** Percentage of CD8+CD4-Cytotoxic T cells, CD8+CD4-T helper cells and CD4+Foxp3+ Treg cells from the spleen. **(P)** Percentage of CD8+CD4+ immature cells and CD8-CD4-DN T cells from the spleen. **(Q)** Percentage of CD69+ Activated cytotoxic T cells from the spleen. **(R)** CD44loCD62L + Naive cytotoxic T cells from the spleen. **(S)** Percentage of CD44hiCD62 – Memory cytotoxic T cells from the spleen. **(T)** Percentage of CD44midCD62 – Effector cytotoxic T cells from the spleen. **(U)** Percentage of CD69+ Activated T helper cells from the spleen. **(V)** CD44loCD62L + Naive T helper cells from the spleen. **(W)** Percentage of CD44hiCD62 + Memory T helper cells from the spleen. **(X)** Percentage of CD44midCD62 – Effector T helper cells from the spleen. Error bars represent SEM.

### Independent Validation Cohort

To ensure the protective effects of β-cell Cre expression were not solely limited to a single animal housing facility, we additionally studied female NOD:*Ins1*^Cre/WT^ mice that were independently generated at a separate site (Fig. 6) in parallel to the cohorts studied in Figure 1. The overall incidence of diabetes development in female NOD mice in this second animal colony was as expected (approximately 65-80% by 25 weeks of age, Fig. 6A). Similar to our data shown in Figure 1, we observed that female NOD:*Ins1*^Cre/WT^ animals were protected from diabetes incidence (25% by 1 year) and had significantly improved mean blood glucose (Fig. 6C) when compared to NOD:*Ins1*^WT/WT^ littermates (75% incidence by 1 year). Again, these studies confirm the protective effects of reduced *Ins1* gene dosage and β-cell Cre expression in NOD mice and suggest that these findings are not due to a consequence of environment or housing.

**Figure 6.**
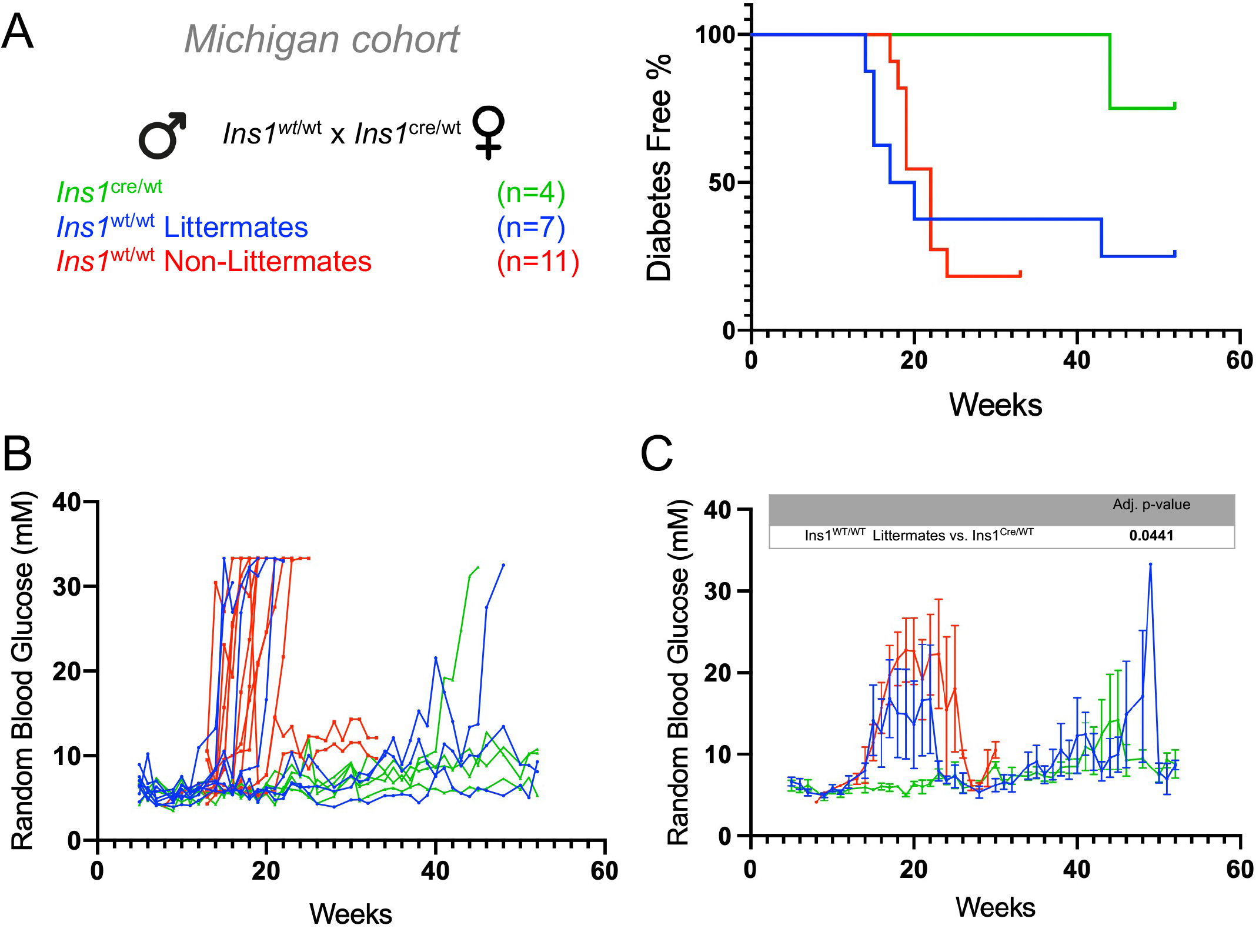
*Ins1* replacement with Cre protects female NOD mice from type 1 diabetes in an independent facility. **(A)** Kaplan-Meier plot denoting diabetes incidence. **(B**,**C)** Individual and mean random blood glucose of female NOD colonies from a second, independent site. The mean blood glucose of the *Ins1*^Neo/Cre^ (green) was significantly lower than that of the *Ins1*^WT/WT^ (blue) littermates, with an adjusted p-value < 0.05. Female NOD non-littermate controls used to track overall diabetes incidence in the colony are listed in red. Error bars represent SEM.

## Discussion

In this study, we used a rigorous littermate control study design to examine the effects, of replacing one *Ins1* allele with Cre-recombinase, on the onset of diabetes in mice of an NOD background. To investigate, whether potential effects were due to the loss of *Ins1* or the introduction of Cre we included *Ins1*^Neo/WT^ mice, in our study. Both *Ins1*^Cre^ and *Ins1*^Neo^ mice have reduced *Ins1* gene dosages with 50% or 100% in their heterozygeous and homozygeous states, respectively. We found a similar reduction in the diabetes incidence in female *Ins1*^Cre/WT^ mice and *Ins1*^Neo/Cre^ mice when compared to littermate control *Ins1*^WT/WT^ mice. This work has implications for our understanding of the pathogenesis of type 1 diabetes, as well as for the use of Cre-recombinase as a tool for *in vivo* genome engineering in mouse models of the disease.

Our findings concur with previous work that showed a similar reduction in diabetes incidence in female NOD mice lacking 1 or 2 alleles of *Ins1* (10,15), although we did not observe additional protection with the double *Ins1* knockout, beyond what was seen with the Cre replacement allele. Similarly, our data are in alignment with a previous study showing replacing the murine *Ins1* gene with the human *INS* gene was found to protect female NOD mice from diabetes in both heterozygous and homozygous states (17). We were unable to detect differences in diabetes incidence in male *Ins1*^Cre/WT^, *Ins1*^Neo/WT^, and *Ins1*^Neo/Cre^ NOD mice, perhaps due to an insufficient study period. A previous study found that removal of a single *Ins1* allele is sufficient to abolish spontaneous diabetes in 50-week-old male NOD mice (10). Together with the work of others, our experiments support the contention that proinsulin 1 is a key player in the generation of autoimmunity in mice. While there are several autoantigens targeted by autoreactive T cells in type 1 diabetes (2), insulin and proinsulin are particularly common autoantibody targets in prediabetic humans (29,30). Our experiments were underpowered to detect subtle differences in the levels of insulin autoantibodies, as we were limited by only examining a single, late time point. However, we did examine insulitis at two time points, and insulin antibodies are often correlated with insulitis (31). In our hands, there was a qualitative difference in the number of islets that did not exhibit insulitis at 50 weeks in mice with at least 1 *Ins1* allele replaced. At 12 weeks, there was a slight trend towards more insulitis-free islets in the *Ins1*^Neo/Cre^ mice, consistent with the greater protection from type 1 diabetes incidence. These observations are consistent with previous studies showing that *Ins1* knockout in NOD mice is protective against the development and severity of insulitis (10). Thus, our results show that reducing the *Ins1* gene dosage lowers the threshold required for diabetes onset, likely by removing the source of primary autoantigens and suppressing insulitis.

An important observation of our study is that Cre expression itself has protective effects in NOD mice, beyond the protection afforded by the loss of 1 *Ins1* allele. To examine the specific consequences of Cre expression, we compared NOD;*Ins1*^Cre/WT^ to a different knock-in (neo) *Ins1*^Neo/WT^ mouse line and found roughly twice as much final diabetes protection in Ins*1*^Cre/WT^ mice versus *Ins1*^Neo/WT^ mice. Though it is possible diabetes could have been delayed further than 1 year, these findings suggest that Cre itself may affect diabetes rates in female NOD mice, with the caveat the Neo and Cre insertions are different, altering the local gene structure. Mechanistically, we are able to attribute the observed effects to differences in insulitis, suggesting the possibility of a β-cell autonomous effect of Cre expression. Previous studies highlighted the potential of Cre recombinase to result in toxicity due to DNA damage (32,33) Mammalian genomes contain pseudo loxP sites, and even though these sequences can deviate considerably from the consensus loxP site, they can still serve as functional recognition sites for Cre (32). It is predicted that the frequency of pseudo-loxP sites could be as many as 250 and 300 in mouse and human genomes, respectively (33). The sustained presence of high levels of Cre in fibroblasts can cause growth arrest and chromosomal abnormalities (18-20,22,23). Cre-dependent DNA damage and accumulation of cytoplasmic DNA have been shown to initiate a STING-dependent immune response (34). STING is an intracellular adaptor molecule, associated with the endoplasmic reticulum membrane (35), that can play a critical role in detecting pathogen-derived DNA in the cytoplasm (36). There is precedence for diabetes protection in NOD mice with early exposure to pathogen in the coxsackievirus mode (37). A recent paper reported that STING is required for normal β-cell function in mice (38), although, ironically, the study did not employ Cre-only controls. Theoretically, Cre expressed in β-cells could delay the onset of diabetes in a STING-dependent manner. Future studies, beyond the scope of this work, will be required to delineate the molecular mechanisms by which Cre expression induces further protection that *Ins1* loss in the NOD mouse model. To unequivocally demonstrate an effect of Cre activity on progression to T1D/insulitis, mice would have to be generated with enzyme-dead Cre knocked into the same locus, but such studies are also beyond the scope of this work.

As with all studies, this work has a number of limitations. For example, the broadly observed phenomenon that diabetes incidences differ between NOD mouse colonies means that we cannot directly compare the results between sites. We do not know the reasons for the apparent difference in diabetes incidence in wildtype NOD mice between our colonies, but it could be related to many environmental factors, including native microbiome, water, food, or bedding, as well as subtle differences in genotype. Another caveat of our study is that we could not simultaneously address *Ins1* gene dosage and the effects of the Cre transgene, because generating both homozygous and heterozygous littermates with each of the knock-in alleles is impossible. In order to reduce the potential impacts of environment and genotype, mentioned above, it was imperative that we prioritized using littermates. Another limitation is the natural uncertainty if *Ins1*^Cre/WT^ mice could have proceeded to develop diabetes beyond the end of this study. While our study has limitations, we believe it is important to report these results that will help guide those in the field who use type 1 diabetes mouse models or Cre in any context.

In summary, our observations suggest caution when interpreting experiments that involve Cre recombinase in NOD mice. Our data showed that Cre expression itself has protective effects in NOD mice, beyond the protection afforded by the loss of 1 *Ins1* allele. Cre-loxP systems are vital tools for research, however, there are multiple caveats that should be considered related to off-target effects and the determination of correct controls. At the bare minimum, Cre-only controls are essential. Additional tools for *in vivo* genome engineering are required to advance the field. Many studies will need to be re-interpreted.

## Acknowledgements

We thank colleagues in the β-cell biology field for helpful discussions, including on Twitter.

## Acknowledgements

We thank the BC Diabetes Research community and the Johnson laboratory members for valuable feedback and discussions at local meetings. We thank Dr. Liping Yu and his team at the Insulin Antibody Core Lab at the University of Colorado, Barbara Davis Diabetes Centre for insulin auto-antibody analysis. We thank Dr. Cara Ellis, at University of Alberta for helpful statistics discussions and analysis of survival/ diabetes incidence data. We thank the amazing animal care services staff for their daily attention and care of our NOD mice housed at UBC, especially during the challenging COVID-19 pandemic.

## Data Availability

Data generated during and/or analyzed during the current study are not publicly available but are available from the corresponding author on reasonable request.

